# New chromosome-level haplotyped genome assemblies and annotation for the Japanese Quail (*Coturnix Japonica*)

**DOI:** 10.64898/2026.05.12.724545

**Authors:** Cédric Cabau, Fabien Degalez, Sophie Leroux, David Gourichon, Rémy-Felix Serre, Caroline Vernette, Cécile Donnadieu, Carole Iampietro, Céline Vandecasteele, Frédérique Pitel, Christophe Klopp

## Abstract

The Japanese quail (*Coturnix japonica*) is a widely used model organism in developmental biology, genetics, and agriculture. Here, we present new, haplotyped, high-quality genome assemblies of the Japanese quail, generated using a combination of state-of-the-art sequencing technologies, including PacBio HiFi long reads, Oxford Nanopore sequencing, and Hi-C scaffolding. This assembly has a total length of 1.19 Gb, 80% of which is included in chromosomes, and is highly complete (BUSCO score aves_odb10: 97.3). Assembly metrics show a marked improvement in contiguity, with a significantly higher scaffold N50 and a lower number of contigs compared to the reference genome assembly. Remarkably, the assembly extends previously truncated chromosome ends, with 31 telomeres detected. In addition, we merged the existing Ensembl and Refseq annotations and obtained a combined set of 26,102 genes, of which 25,038 genes were successfully mapped on the improved assembly haplotype 1 (Cjap1.hap1). Together, these new genome assemblies and their enriched annotation provide a robust genomic framework for future research. They enhance our ability to investigate developmental processes, genetic and epigenetic inheritance, and host-pathogen interactions. Furthermore, they offer valuable insights for conservation genetics and sustainable breeding programs. This resource represents a critical step forward in leveraging the full potential of the Japanese quail as a model species in both basic and applied research.

## Introduction

The Japanese quail (*Coturnix japonica*) is an avian species of significant biological, ecological, and economic importance. As a member of the family Phasianidae within the order Galliformes, it is closely related to chicken (*Gallus gallus*), another widely studied species.

Thanks to its rapid reproductive cycle and small size, which facilitate laboratory rearing, the Japanese quail has long served as a widely used model organism across numerous disciplines. It has been particularly influential in developmental biology through the pioneering quail-chick chimera technique (Le Douarin 1973). This method has enabled transformative insights into embryonic development by offering unparalleled access to embryos (Le Douarin 2005). Japanese quail has also been extensively used in research spanning physiology (Balthazart et al. 2003), genetics (Kayang et al. 2006), behavior (Mignon-Grasteau et al. 2003), or biomedicine (Serralbo et al. 2020; Lin et al. 2002). The generation of transgenic quail lines has further expanded its utility, providing invaluable tools for cutting-edge biological and biomedical investigations (Barzilai-Tutsch et al. 2022; Huss et al. 2015; Serralbo et al. 2020).

Among farm animals, the Japanese quail stands out as a model of choice for multi/transgenerational studies. Its low maintenance costs, short reproductive cycle, and the feasibility of environmental manipulations during embryogenesis make it ideal for exploring genetic and epigenetic inheritance mechanisms (Leroux et al. 2017; Vitorino Carvalho et al. 2023). In addition to its prominent role as a research model, the Japanese quail holds considerable agricultural value. Renowned for its flavorful eggs and meat, it is farmed worldwide, ranking second only to the domestic chicken in poultry production for meat and eggs (Lukanov 2019). While domesticated quail populations thrive in agricultural systems, wild populations of the closely related common quail (*Coturnix coturnix*) face increasing threats due to habitat loss and disruptions of migratory networks (Sáez et al. 2023). The hybridization between these two taxa in the wild poses a challenge for conserving the genetic integrity of the common quail, emphasizing the need for targeted biodiversity preservation efforts (Chazara et al. 2010); (Sanchez-Donoso et al. 2016).

The presence of numerous tiny, repeat- and GC-rich microchromosomes poses a significant challenge for the assembly of bird genomes, and these are often underrepresented or entirely missing from draft assemblies (Peona et al. 2018). A previous genome assembly for the Japanese quail has been published (Morris et al. 2020), underscoring the utility of high-quality genomic resources for addressing scientific questions related to social behavior, seasonal biology, and host-pathogen interactions, including responses to H5N1 influenza infection. While the previously published assembly was comparable in quality to the chicken genome assembly available at that time, recent advancements in sequencing and assembly technologies have since enabled significant improvements. The latest chicken genome reference version (GRCg7b; Smith et al. 2023) exemplifies these advances, providing an exceptionally high-quality reference.

Building on these technological advances, we employed state-of-the-art methods, including high quality long-read sequencing, to generate an improved genome assembly for the Japanese quail. In addition, we merged the existing Ensembl and Refseq annotations to annotate the new assembly.

This enhanced genome assembly and its annotation represent an important resource for advancing our understanding of evolution, ecology, and adaptation in Galliformes. It also paves the way for practical applications in conservation biology and agriculture.

## Material and methods

### Biological material and DNA sequencing

The ONT and PacBio HiFi data used in this study have been described in Terzian et al, 2025 (Terzian et al. 2025).

Briefly, blood DNA was extracted from a 6 month-old female quail of a high social reinstatement line (Mills et Faure 1991), using a high-salt extraction method (Roussot et al. 2003).The animal was bred at INRAE, UE1295 Pôle d’Expérimentation Avicole de Tours, F-37380 Nouzilly in accordance with European Union Guidelines for animal care, following the Council Directives 98/58/EC and 86/609/EEC. The facility is registered with the French Ministry of Agriculture under license number C37–175–1 for animal experimentation (https://doi.org/10.15454/1.5572326250887292E12).

ONT libraries were prepared following the manufacturer’s instructions and sequenced on a PromethION instrument (Oxford Nanopore Technologies, UK). PacBio library preparation was performed according to the manufacturer’s instructions and sequenced by diffusion loading onto 2 SMRT cells on a Sequel II instrument (PacBio, USA).

The Hi-C data were obtained from the same blood sample. The Hi-C library was generated as in (Du et al. 2020), according to a protocol adapted from (Foissac et al. 2019), and amplified using paired-end primers (Illumina) for 10 cycles. The sequencing generated a total of 106 million paired reads (2 × 150 bp), produced with Illumina NovaSeq 6000 and HiSeq 3000.

### Genome assembly, validation and scaffolding in chromosomes

Three read types including HiFi, ONT long and Hi-C reads were used to produce contigs with Hifiasm version 0.24.0 (Cheng et al. 2021). The corresponding reads accessions are ERR11528759 and ERR11528760 for HiFi, ERR11591487 and ERR11591488 for ONT and ERR13583727 and ERR13583728 for Hi-C. Two haplotyped contigs sets were produced corresponding to hap1 and hap2. Both contigs sets were processed with MitoHiFi (Uliano-Silva et al. 2023) to find the mitochondrion contigs.

Hap1 was scaffolded with chromosome conformation Hi-C data. These data were processed with the juicer pipeline v1.6 (Durand, Shamim, et al. 2016). The assembly was scaffolded with 3d-dna v180419 (Dudchenko et al. 2017) and manually corrected with juicebox v2.13.05 (Durand, Robinson, et al. 2016). Hap2 was scaffolded with RagTag v2.1.0 (Alonge et al. 2022) using scaffolded hap1 as reference. All scaffolds were evaluated and checked for contamination using blobToolKit version 4.4.2 (Challis et al. 2020) with default parameters. Assembly metrics were produced with the assemblathon_stats.pl (https://github.com/KorfLab/Assemblathon/blob/master/assemblathon_stats.pl) script. The scaffold protein content was checked with BUSCO version 5.7.1 (Manni et al. 2021) in euk_genome_aug mode using the aves_odb10 lineage and default parameters. The assembly metrics and busco scores were compared with the public reference genome assembly Coturnix_japonica_2.1 (GCA_001577835.2) To evaluate the extent of the assembly, we searched for telomeric repeats at the ends of each chromosome. We plotted the density of TTAGGG repeats with karyoploteR v1.24.0 (Gel et Serra 2017) using 200 kb windows and a maximum value of 500. The presence of a telomere is indicated by a red star at a chromosome end if at least 100 repeated motifs have been located in the first or last 10 kb.

To describe the enhancement in chromosome continuity, unlocalized contigs from each chromosome of the GCA_001577835.2 reference assembly were aligned with minimap v2.2.28 against the corresponding chromosome in the new Cjap1.hap1 assembly. For an unlocalized contig of the reference assembly to be considered correctly positioned in a scaffold of the new Cjap1.hap1 assembly, the observed match must have the maximum mapping quality (60) and cover at least 90% of the contig length.

### Initial reference assemblies and annotation (Coturnix_Japonica_2.0 & 2.1)

For Refseq, we used the Coturnix_japonica_2.1 assembly (GCF_001577835.2), corresponding to the GenBank assembly GCA_001577835.2. This version includes the mitochondrial genome and its associated annotation from Refseq Release 101. The genomic sequence and gene annotation were downloaded directly from NCBI’s FTP repository (https://ftp.ncbi.nlm.nih.gov/genomes/all/GCF/001/577/835/GCF_001577835.2_Coturnix_japonica_2.1/GCF_001577835.2_Coturnix_japonica_2.1_genomic.fna.gz & https://ftp.ncbi.nlm.nih.gov/genomes/all/GCF/001/577/835/GCF_001577835.2_Coturnix_japonica_2.1/GCF_001577835.2_Coturnix_japonica_2.1_genomic.gtf.gz). Ensembl Release 114 is based on *Coturnix japonica* 2.0 (GCA_001577835.1), a previous version of the same genome.

To ensure consistency across data sources, we harmonized the reference assemblies and annotations used by Refseq and Ensembl, which rely on different versions of the Coturnix japonica genome: according to the NCBI Assembly Report (https://www.ncbi.nlm.nih.gov/datasets/genome/GCF_001577835.2), the only differences between versions 2.0 and 2.1 concern scaffolds LSZS01008902 and KQ966558 which were moved from chromosome W to “unplaced”, and the W chromosome (GenBank record CM003811.1) which was removed from the assembly. These corrections reflect the biological fact that the sequenced individual was a male and therefore lacked a W chromosome. We retrieved the Ensembl annotation based on version 2.0 and manually inspected genes assigned to chromosome W. We found that all genes previously assigned to W in the Ensembl annotation mapped to scaffold NW_015438892.1 in the Refseq assembly. To ensure compatibility with the corrected assembly and avoid sex chromosome artifacts, we excluded these five genes (including nine transcripts) annotated on the W chromosome in the Ensembl annotation.

### Formatting of Refseq and Ensembl genome annotations

Sequence names across both Refseq and Ensembl annotations were standardized to follow Ensembl naming conventions. Assembled chromosomes were labeled using their canonical chromosome names, while unlocalized and unplaced scaffolds retained their GenBank accession numbers. GTF attribute fields were formatted to align with Ensembl specifications. For the Refseq annotation in particular, we constructed gene identifiers by concatenating the “LOC” tag with the corresponding *db_xref* value from the attribute column. In cases where transcripts lacked an associated gene model most commonly on the mitochondrial chromosome, where Refseq used the “unassigned_gene” tag, we generated synthetic gene models by grouping transcripts based on their genomic coordinates. Each of these gene models spanned the minimum start to maximum end positions of the corresponding transcripts, and we assigned a unique gene identifier in the format “LOCfkUnGene0[n]”. We also redefined exon identifiers within Refseq using the structure: “RSEX” followed by the transcript identifier, a separator “0”, and the exon number (*e.g.,* RSEX01587912701). For genes not associated with any transcript, typically pseudogenes, we did not generate synthetic transcript models.

### Merging Refseq and Ensembl genome annotations

To preserve overlapping or nested genes of different biotypes, Refseq and Ensembl annotations were first partitioned by size and biotype before merging. This resulted in: (i) long genes, including protein-coding genes (PCGs), long non-coding RNAs (lncRNAs), and pseudogenes, and (ii) short genes, including small biotypes such as miRNA, rRNA, tRNA, scaRNA, snoRNA, snRNA, and sRNA.

We performed the annotation merge at the exonic chain level and reformatting using two modules from the AGAT toolkit (Dainat et al. 2025): *agat_sp_merge_annotations.pl* to combine the annotations across sources, and *agat_convert_sp_gff2gtf.p*l to convert the resulting GFF3 file into GTF format. To maintain structural and semantic consistency, we retained the original subfields in the GTF attribute column and added additional fields to preserve source-specific information. At the gene level, the fields *previous_gene_id*, *previous_gene_name* and *previous_gene_biotype* record unique identifiers, names and biotypes from either Refseq or Ensembl when a collapse occurred during the merge. At the transcript level, we retained source-specific gene and transcript identifiers, along with the corresponding gene biotype. When merging transcripts with identical exon structures, we assigned the IDs of the removed transcripts to the *equivalent_transcript* field in the merged annotation. This strategy preserved the integrity of original source data while resolving redundancy and enabling downstream comparative and functional analyses.

**Transfer of the enhanced genome annotation from Coturnix_japonica_2.1 to Cjap1.hap1**

We transferred the enhanced annotation to the new *Coturnix japonica* genome assembly (Cjap1.hap1) using the LiftOff pipeline (Shumate and Salzberg 2021), which relies on minimap2 (Li 2018) for spliced alignment. The mapping was performed with the following parameters: *-a --end-bonus 5 --eqx -N 50 -p 0.5*. To correct artifacts introduced during lift-over, we enabled the *--polish* option. This feature realigns the exons and reconstructs coding sequences to preserve open reading frames and transcript integrity wherever possible.

In addition to lifted coordinates, two quality metrics for each gene model were computed: coverage, defined as the proportion of the reference gene’s length aligned to the target genome, and sequence identity, defined as the percentage of aligned nucleotides that remain unchanged between the reference and lifted annotations.

### Generation of chain files and chromosome assignment in the assembly

Chain files allowing coordinate conversion from the Coturnix_japonica_2.1 reference assembly to the new Cjap1.hap1 assembly were generated using the nf-lo pipeline (Talenti et Prendergast 2021). The alignment was carried out using the *--aligner blat* option, which is recommended for same-species lift-overs, along with the *--distance balanced* setting to optimize the trade-off between alignment precision and computational efficiency while conforming to UCSC standards. This configuration was chosen based on benchmarking data from same-species alignments involving the human reference genomes GRCh38 and GRCh37. In parallel, a second set of chain files was generated to enable coordinate conversion from the chicken (Gallus gallus) GRCg7b reference assembly to Cjap1.hap1, representing an estimated divergence of approximately 39 million years. For this cross-species alignment, the *--aligner lastz* option was selected, as recommended for more divergent genomes, in combination with the *--distance medium* setting. The chain file generated between both quail assemblies was first employed to assign chromosomal identities to the new *Cjap1.hap1* scaffolds, focusing on the 10 macrochromosomes (1 to 10), the 19 previously annotated microchromosomes (11 to 28), and the Z sexual chromosome. However, due to the larger number of annotated microchromosomes in the chicken genome (30 assigned), the chicken-to-quail chain file was subsequently used to refine microchromosome assignments.

### Biological samples used for gene expression analysis

We analyzed gene expression using a publicly available RNA-seq dataset comprising seven tissues (https://pmc.ncbi.nlm.nih.gov/articles/PMC7017630/): brain, heart, intestine, kidney, liver, lung, and muscle. Each tissue was collected from a single individual, with six biological replicates per tissue. This dataset is available under the NCBI BioProject accession PRJNA296888.

### Gene expression quantification and expression criteria

We quantified gene expression by aligning RNA-seq reads to the newly assembled *Coturnix japonica* assembly (Cjap1.hap1) using the corresponding lifted-enhanced GTF annotation. Read alignment and quantification were carried out with the nf-core/rnaseqpipeline (v3.16.0) (Patel et al. 2024), employing STAR as the aligner and RSEM for transcript quantification (*--aligner star_rsem*). The pipeline produced both raw read counts and TPM (transcripts per million) normalized expression values for each gene and transcript.

### Tissue-specificity analysis

The tau (τ) metric was used (Yanai et al. 2005), providing a score between 0 (gene expressed at the same level in all tissues) and 1 (gene expressed in exactly one tissue) and following the formula:

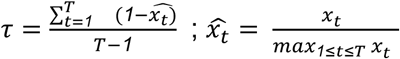

with *x_t_* the expression of the gene of interest in the tissue *t* and among the *T* tissues.

## Results

### Overview of the genome assembly and scaffolding in chromosomes

A total of 1.88 million HiFi reads, 4.15 million ONT reads and 212.65 million Hi-C reads corresponding to 30.54 Gb, 65.00 Gb and 31.90 Gb respectively were assembled in 751 contigs (Table 1).

**Table 1.** Statistics of the genome assembly and scaffolding in chromosomes between Coturnix_japonica_2.1 public reference and Cjap1.hap1 novel assembly. Statistics were produced with the assemblathon_stats.pl script.

|  | GCF_001577835.2 |  | Cjap1.hap1 |  |
| --- | --- | --- | --- | --- |
|  | contigs | scaffolds | contigs | scaffolds |
| Number of contigs/scaffolds | 8,466 | 2,012 | 751 | 376 |
| Number of contigs in scaffolds | 7,163 |  | 400 |  |
| Number of contigs not in scaffolds | 1,303 |  | 351 |  |
| Total size of contigs/scaffolds | 917,278,609 | 927,656,957 | 1,191,131,709 | 1,191,319,209 |
| Longest contig/scaffold | 8,486,926 | 175,656,249 | 31,684,377 | 179,650,266 |
| Shortest contig/scaffold | 58 | 237 | 15,452 | 15,452 |
| Number of contigs/scaffolds > 1K nt | 7,911 | 1,991 | 751 | 376 |
| Number of contigs/scaffolds > 10K nt | 2,776 | 621 | 751 | 376 |
| Number of contigs/scaffolds > 100K nt | 1,519 | 65 | 533 | 261 |
| Number of contigs/scaffolds > 1M nt | 222 | 29 | 214 | 95 |
| Number of contigs/scaffolds > 10M nt | 0 | 17 | 27 | 18 |
| Mean contig/scaffold size | 108,349 | 461,062 | 1,586,061 | 3,168,402 |
| Median contig/scaffold size | 4 151 | 4,462 | 294,398 | 234,283 |
| N50 contig/scaffold length | 794,327 | 82,189,922 | 6,796,293 | 55,622,978 |
| L50 contig/scaffold count | 315 | 4 | 46 | 6 |
| contig/scaffold %A | 29.34 | 29.01 | 28.61 | 28.61 |
| contig/scaffold %C | 20.62 | 20.39 | 21.29 | 21.29 |
| contig/scaffold %G | 20.66 | 20.43 | 21.44 | 21.43 |
| contig/scaffold %T | 29.39 | 29.06 | 28.66 | 28.66 |
| contig/scaffold %N | 0.00 | 1.12 | 0.00 | 0.02 |

A set of 811.75 million reads was generated by combining Hi-C sequencing with a publicly available Hi-C run (SRR24088247), 98.35% of which were alignable to the assembled contigs. Starting from these, 169.31 million chromatin contacts were identified, distributed across inter-and intra-chromosomal links with 60.71 and 108.60 million contacts, respectively. Of the intra-chromosomal contacts, 32% were found to be larger than 20 Kb. Following scaffolding manual curation, a chromosome-scale assembly of 1.19 Gb in 346 scaffolds was built. The snail plot in Figure 1 provides a summary of the assembly statistics, indicating the distribution of scaffold lengths and other assembly metrics. The distribution of assembly scaffolds on GC proportion and coverage is shown in Figure 2. Figure 3 presents a cumulative assembly plot, with curves for subsets of scaffolds assigned to different phyla, illustrating the completeness of the assembly.

**Figure 1.**
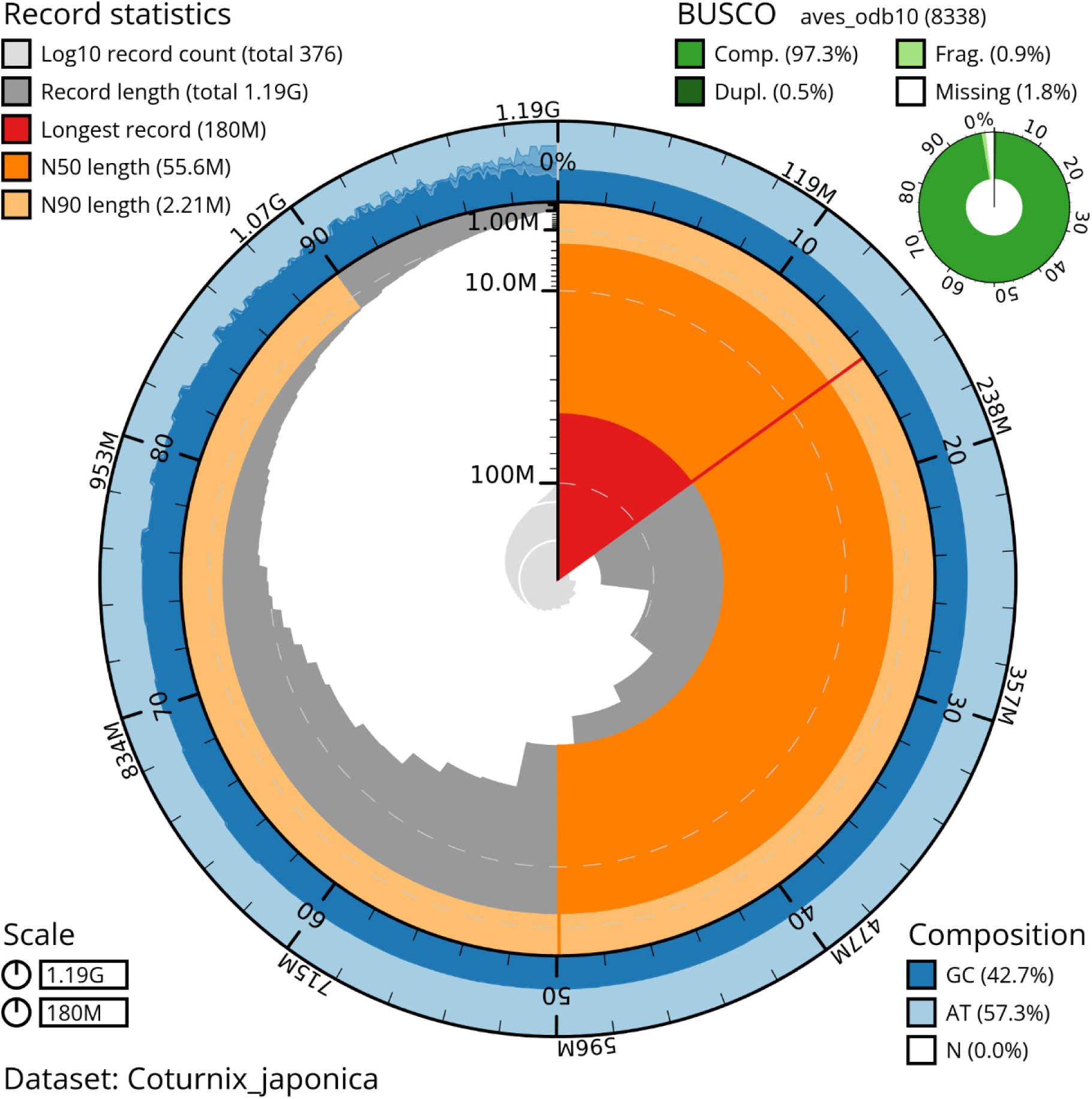
Genome assembly of Coturnix japonica, Cjap1.hap1: metrics. The BlobToolKit snail plot shows N50 metrics and BUSCO gene completeness. The main plot is divided into 1,000 bins around the circumference with each bin representing 0.1% of the 1,191,319,209 bp assembly. The distribution of scaffold lengths is shown in dark grey with the plot radius scaled to the longest scaffold present in the assembly (179,650,266 bp, shown in red). Orange and pale-orange arcs show the N50 and N90 scaffold lengths (55,622,978 and 2,207,750 bp), respectively. The pale grey spiral shows the cumulative scaffold count on a log scale with white scale lines showing successive orders of magnitude. The blue and pale-blue area around the outside of the plot shows the distribution of GC, AT and N percentages in the same bins as the inner plot. A summary of complete, fragmented, duplicated and missing BUSCO genes in the aves_odb10 set is shown in the top right.

**Figure 2.**
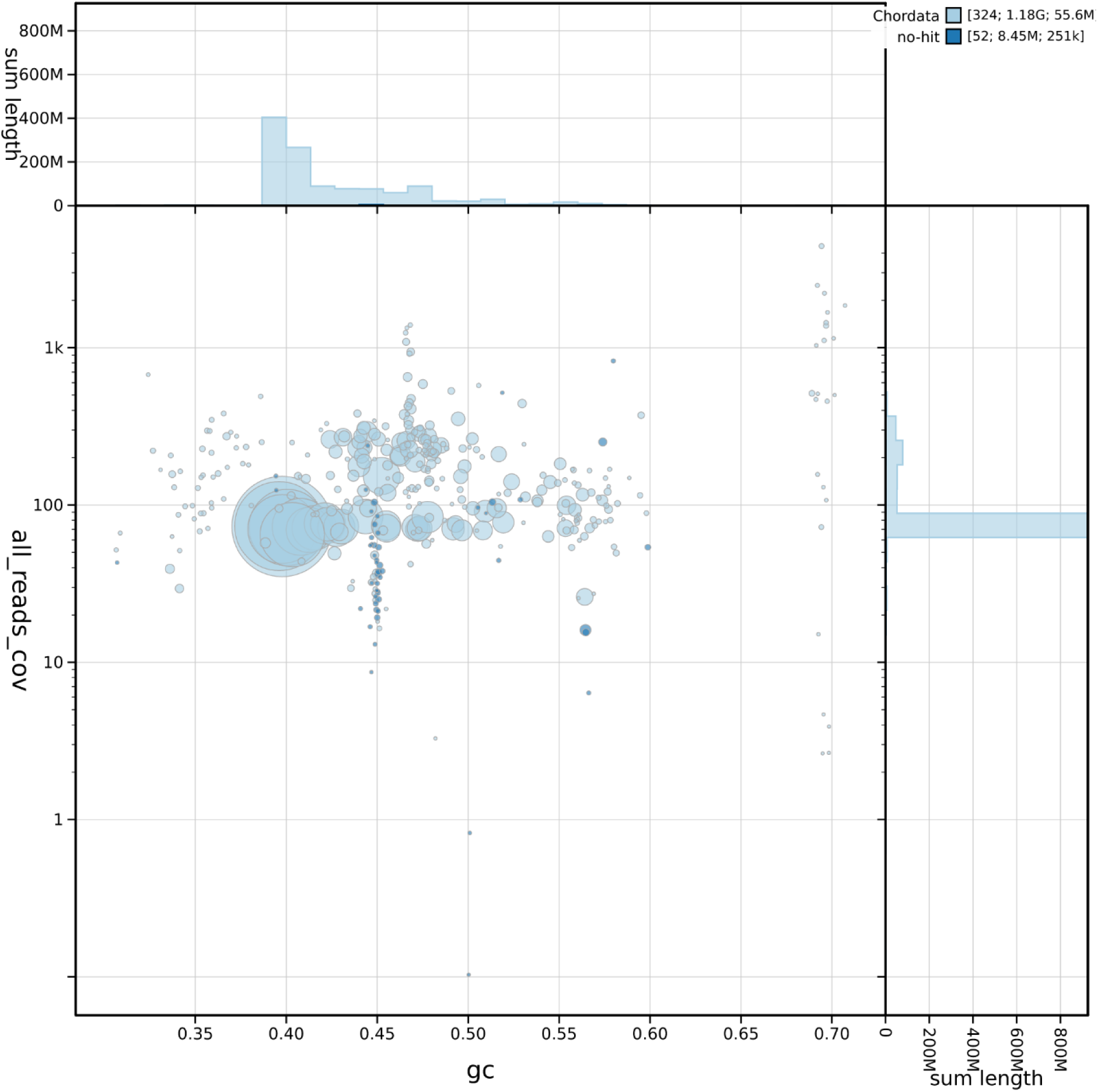
Genome assembly of Coturnix japonica, *Cjap1.hap1*: GC-coverage plot. The circles represent scaffolds, with the size proportional to scaffold length.The histograms along the axes display the total length of sequences distributed across different levels of coverage and GC content. Color of circles indicate taxonomic hits of each phylum represented in the assembly.

**Figure 3.**
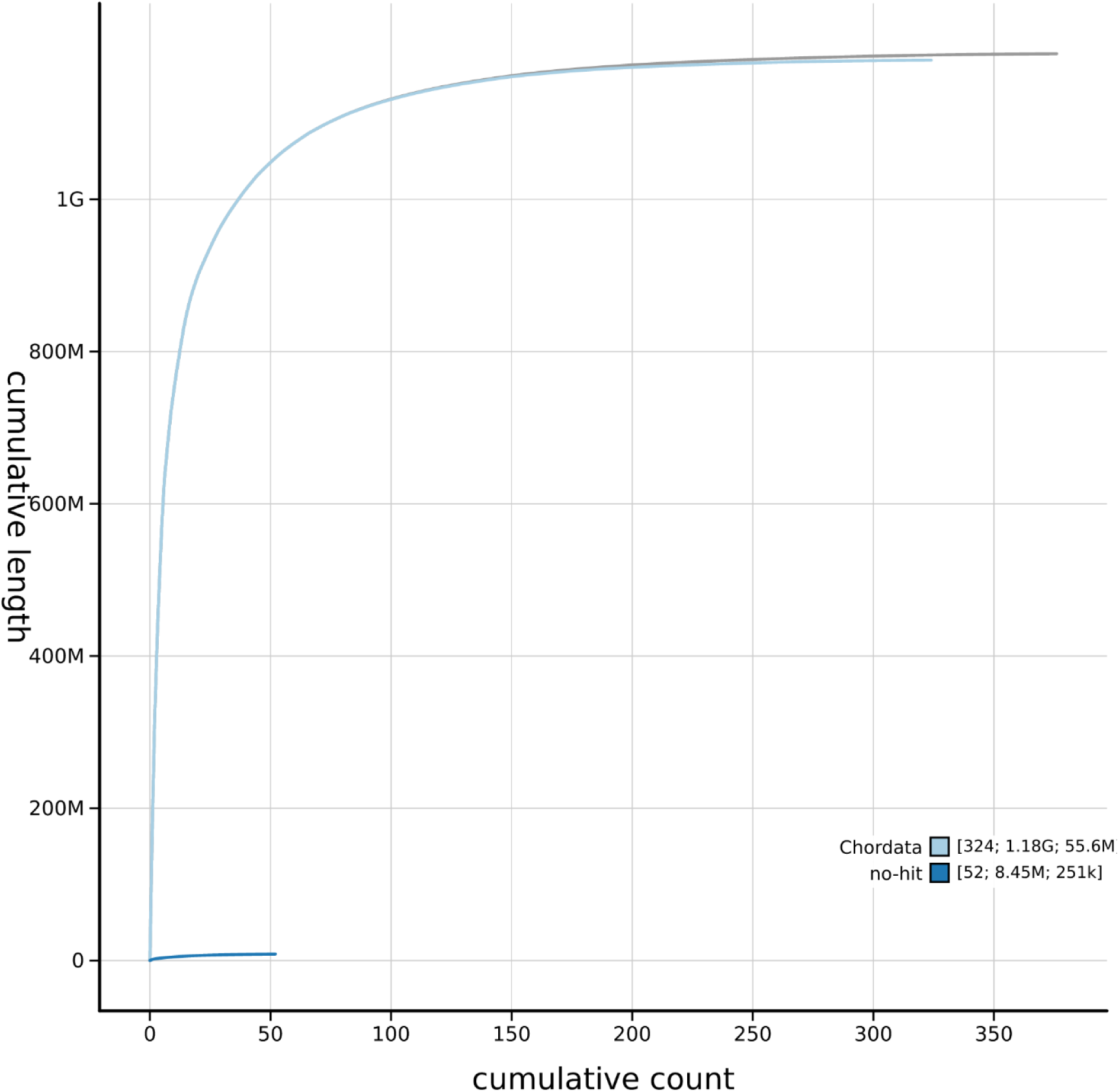
Genome assembly of Coturnix japonica, Cjap1.hap1: cumulative sequence plot. The grey line shows cumulative length for all scaffolds. Coloured lines show cumulative lengths of scaffolds assigned to each phylum using the buscogenes taxrule.

With a total length of 1.19 Gb and 0.95 Gb of the assembly scaffolded into 29 chromosomal pseudomolecules, the final assembly is much larger than the reference assembly (0.93 Gb), adding 50 Mb to chromosomes. The N50 contig length of 6.8 Mb is also much larger compared to 0.79 Mb of the reference. The smaller N50 scaffold length is due to the increased assembly size. Indeed, while the four largest scaffolds in the new assembly are larger than their counterparts in the reference assembly, six of the largest scaffolds are required to reach half of the assembly’s total size. Both assemblies have very close BUSCO scores as presented in Table 2.

**Table 2.** Comparison of BUSCO scores between Coturnix_japonica_2.1 public reference and Cjap1.hap1 novel assembly using aves_odb10 lineage corresponding to 8,338 proteins.

|  | Complete | Single | Duplicate | Fragmented | Missing |
| --- | --- | --- | --- | --- | --- |
| Coturnix_japonica_2.1<br>(GCF_001577835.2) | 96.5 | 96.3 | 0.2 | 0.6 | 2.9 |
| Cjap1.hap1 | 97.3 | 96.8 | 0.5 | 0.9 | 1.8 |

Regarding telomere presence verification, five chromosomes showed telomeric repeats at both ends, 21 chromosomes showed them at one end, and none were detected in the five shortest chromosomes of the assembly (Figure 4). It should be noted that no telomeres were detected in the Coturnix_Japonica_2.1 assembly (Figure 5).

**Figure 4.**
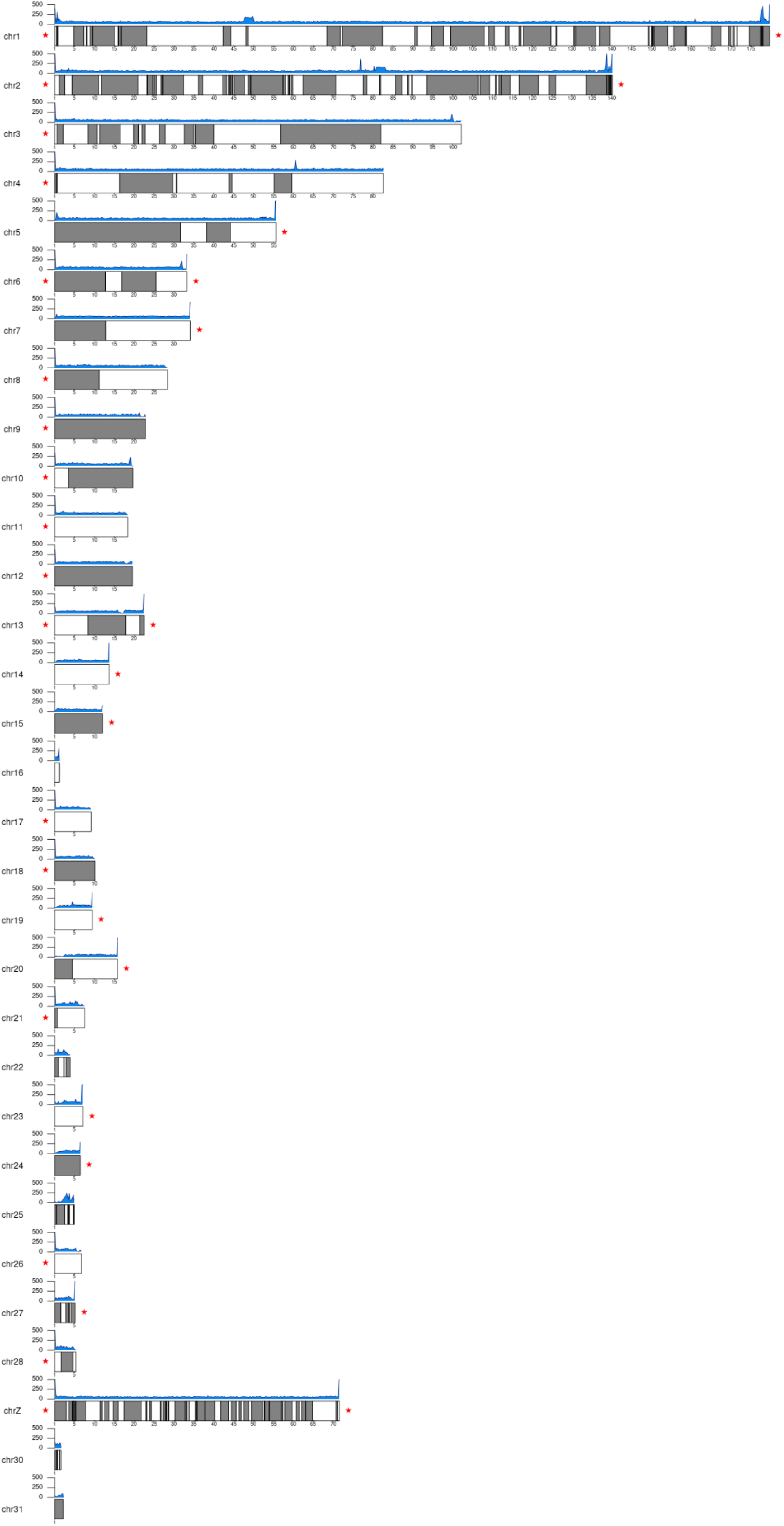
Genome assembly of Coturnix japonica, Cjap1.hap1: telomeric repeats density. The blue areas show the density of TTAGGG repeats along chromosomes in 200 kb windows. Cytoband like coloration Red stars indicate the presence of a telomere at the marked extremities.

**Figure 5.**
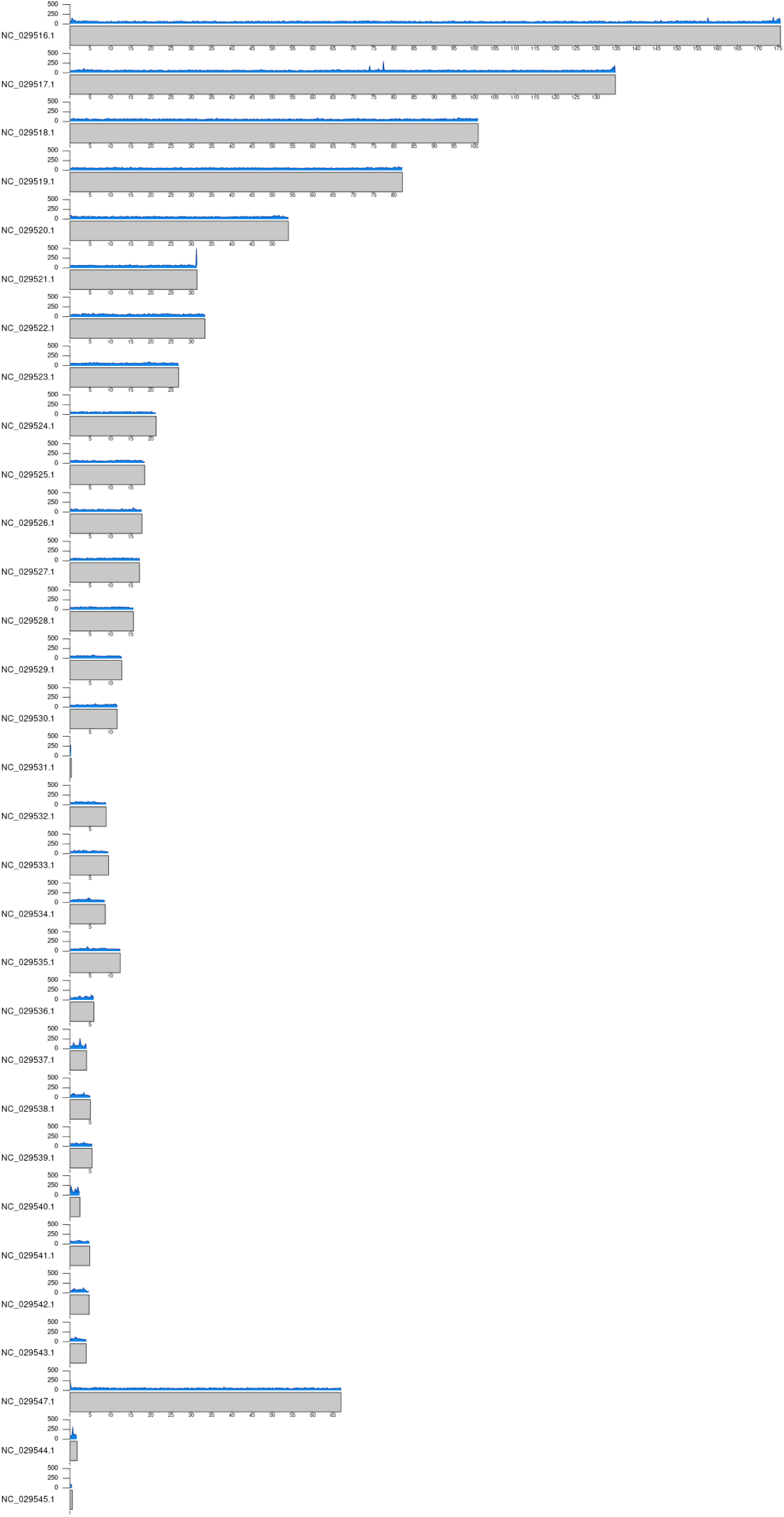
Reference assembly of Coturnix japonica, Coturnix_japonica_2.1: telomeric repeats density. The blue areas show the density of TTAGGG repeats across chromosomes in 200 kb windows.

### Manual assignation of the contigs from the new *Cjap1.hap1* assembly

To evaluate improvements in assembly and chromosome-level resolution, we aligned scaffolds from the new Cjap1.hap1 assembly to both the previous quail (Coturnix_japonica_2.1) and the chicken (*Gallus gallus*, GRCg7b) reference genome assemblies. Chromosome-scale structure was largely conserved: for the 10 previously annotated macrochromosomes and the Z sex chromosome, Coturnix_japonica_2.1 scaffolds aligned almost entirely within their corresponding Cjap1.hap1 chromosomes, indicating strong collinearity (Figure 6A). In addition, the 19 microchromosomes previously annotated in Coturnix_japonica_2.1 were recovered (Figure 6B) and confirmed through alignment with the chicken genome. Of the unlocalized contigs in Coturnix_japonica_2.1 associated with well-characterized chromosomes (1–28 and Z), approximately 75% are now anchored to their respective chromosomes in Cjap1.hap1 (Table 3). Notably, all unlocalized contigs from eight chromosomes were fully placed, representing a substantial improvement in contiguity and chromosomal assignment. For example, chromosome 1 (HiC_scaffold_1 in Cjap1.hap1; NC_029516.1 in Coturnix_japonica_2.1) increased from 175,656,249 bp in Coturnix_japonica_2.1 (with 251 unlocalized contigs totaling 801,044 bp) to 179,650,266 bp in Cjap1.hap1, with 185 of those contigs successfully placed (Figure 6C). This 2.3% size increase likely reflects also the incorporation of previously unplaced contigs (n=23) and unannotated repetitive sequences. Comparative alignment with the chicken genome also supports the putative identification of several additional microchromosomes not annotated in Coturnix_japonica_2.1. For instance, HiC_scaffold_89 (1,142,415 bp), which includes nine previously unplaced contigs, aligns with chicken microchromosome 30 (NC_052561; 755,666 bp), and HiC_scaffold_143 (472,852 bp) aligns with three previously unplaced contigs and chicken microchromosome 38 (NC_052569; 667,312 bp).

**Figure 6.**
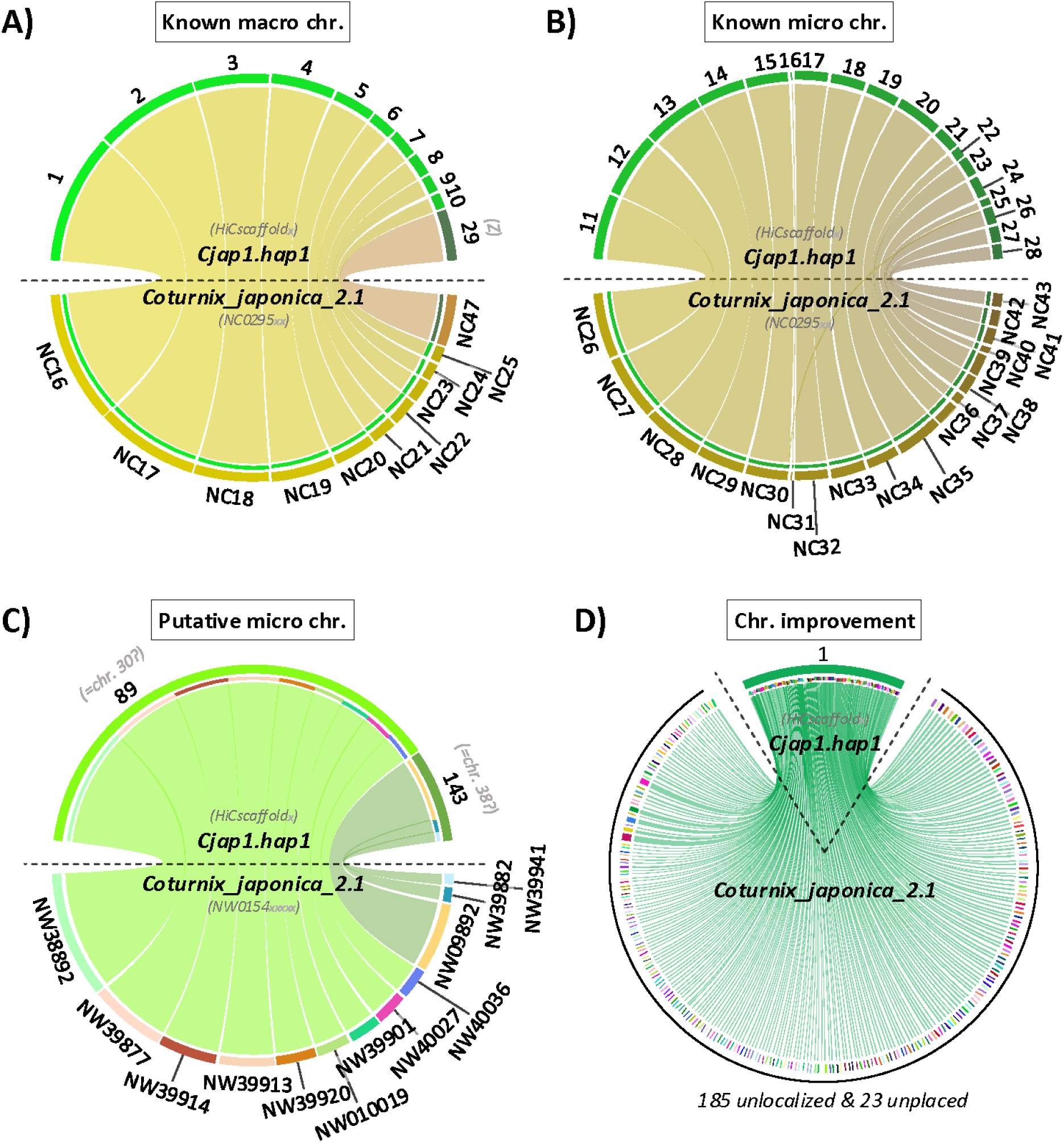
Correspondence between contigs of *Coturnix_japonica_2.1* and the new Cjap1.hap1 assembly. **A,** Previously known macrochromosomes (chr.). **B,** Previously known microchromosomes **C,** Putative new microchromosome incorporating previously unassigned contigs. **D,** Improvement of chromosome 1 through the integration of previously unlocalized and unplaced contigs.

**Table 3.** Number of previously unlocalized contigs from the Coturnix_japonica_2.1 assembly now assigned to well-identified chromosomes in the new Cjap1.hap1 assembly.

| Chromosome | Number of unlocalized contigs in the *Coturnix japonica*_2.1 assembly | Number of well localized contigs in Cjap1.hap1 | % of well localized contigs in Cjap1.hap1 | Number of unplaced contigs in the *Coturnix japonica*_2.1 assembly assigned in Cjap1.hap1 |
| --- | --- | --- | --- | --- |
| *chr1* | 251 | 185 | 74% | 23 |
| *chr2* | 196 | 148 | 76% | 30 |
| *chr3* | 118 | 85 | 72% | 9 |
| *chr4* | 51 | 45 | 88% | 1 |
| *chr5* | 49 | 35 | 71% | 5 |
| *chr6* | 15 | 12 | 80% | 2 |
| *chr7* | 23 | 16 | 70% | 2 |
| *chr8* | 8 | 7 | 88% | 3 |
| *chr9* | 4 | 3 | 75% | 1 |
| *chr10* | 3 | 3 | 100% |  |
| *chr11* | 8 | 6 | 75% | 2 |
| *chr12* | 4 | 2 | 50% | 1 |
| *chr13* | 6 | 5 | 83% | 1 |
| *chr14* | 5 | 4 | 80% |  |
| *chr15* | 3 | 3 | 100% | 1 |
| *chr16* |  |  |  | 1 |
| *chr17* | 3 | 3 | 100% |  |
| *chr18* | 3 | 1 | 33% |  |
| *chr19* | 3 | 3 | 100% |  |
| *chr20* | 4 | 3 | 75% |  |
| *chr21* | 1 | 1 | 100% | 1 |
| *chr22* |  |  |  | 2 |
| *chr23* | 3 | 2 | 67% | 1 |
| *chr24* | 3 | 3 | 100% | 2 |
| *chr25* | 2 | 2 | 100% | 5 |
| *chr26* | 1 | 0 | 0% | 2 |
| *chr27* | 4 | 2 | 50% | 2 |
| *chr28* | 1 | 1 | 100% | 2 |
| *chrZ* | 108 | 83 | 77% | 10 |
| **all** | **880** | **663** | **75%** | **109** |

### Merging of Ensembl and Refseq annotations for the Coturnix_japonica_2.1 assembly

A total of 15,729 protein-coding genes (PCGs) were annotated by Ensembl, while 16,026 PCGs were identified in the Refseq annotation (Figure 7A). For long non-coding RNAs (lncRNAs), 5,086 gene models were reported in the Ensembl annotation, compared to 4,222 in Refseq. In total, 21,554 genes were annotated by Ensembl and 20,980 by Refseq. To create a more comprehensive annotation resource, both datasets were merged using the AGAT tool, with gene models divided into two biotype categories (short/long) as described in the Materials and Methods. Through this process, a combined set of 26,102 genes was finally annotated, including 16,761 PCGs and 8,171 lncRNAs. This represents an increase of more than 17% in total gene models relative to the annotations considered alone. The most pronounced gain, however, was observed at the transcript level: 86,845 transcripts are now annotated in the merged annotation (68,615 for PCG and 14,774 for lncRNA), compared to 37,593 in Ensembl and 49,534 in Refseq (Figure 7A). However, when transcript models were merged based on intron-chain equivalence, the total number of transcripts decreased to 79,679 corresponding to a reduction of 8.3% reflecting that even if some divergences are observed only for UTR between isoforms, a majority of exonic parts are different between annotations.

**Figure 7.**
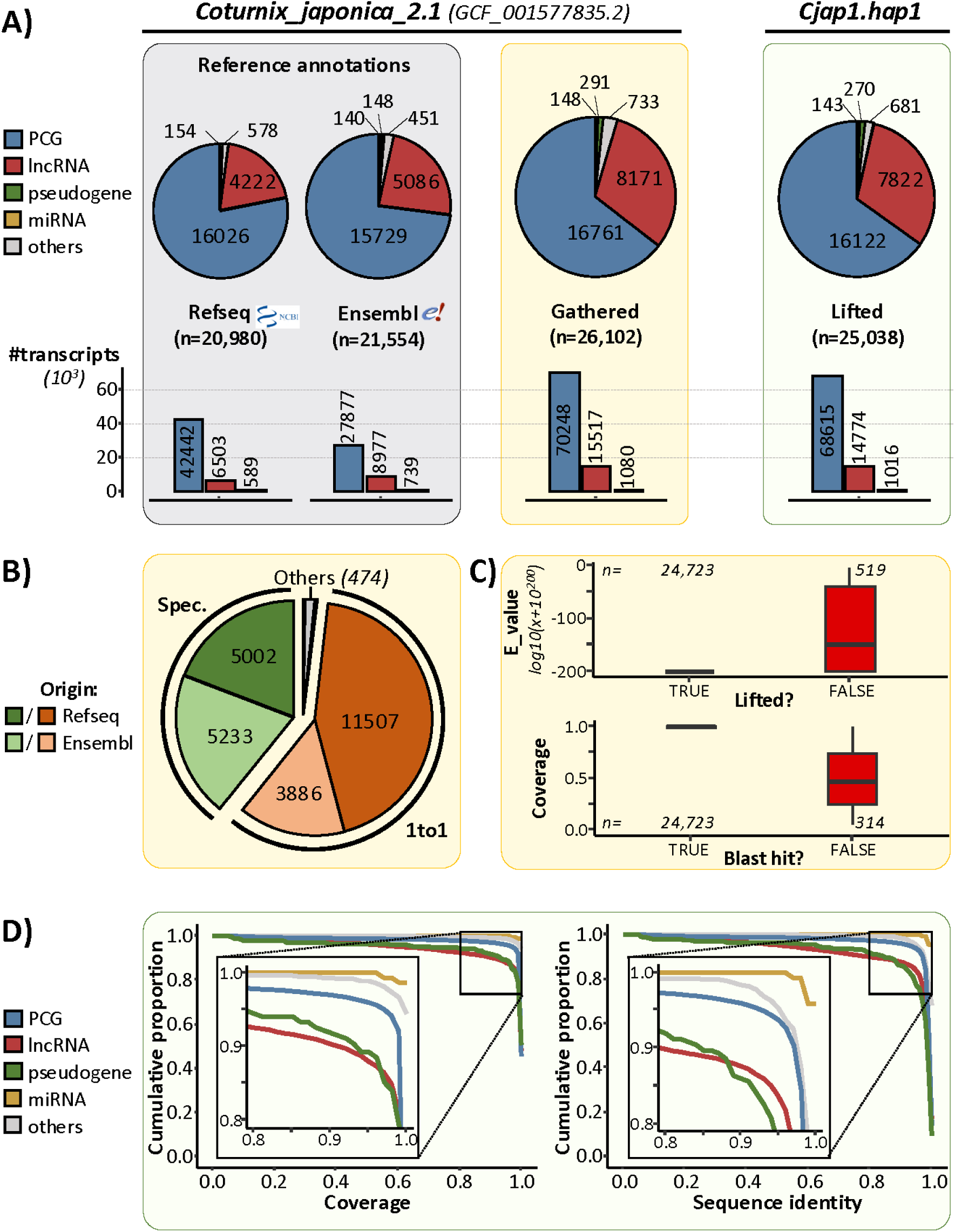
Improved gene annotation in the Cjap1.hap1 assembly. **A,** Number of genes and transcripts by biotype in Refseq and Ensembl reference annotations (grey), the merged annotation (yellow), and the lifted annotation mapped to the new Cjap1.hap1 assembly (green). **B,** Overlap and unique gene models between Refseq and Ensembl annotations. **C,** Distribution of BLASTN E-values for lifted genes in the enhanced annotation, and coverage relative to the presence of a BLAST hit in the lifted set. **D,** Coverage and sequence identity of lifted genes by biotype.

In detail concerning overlap between annotations, a total of 15,393 genes were found to be shared between the two sources with a strict one-to-one correspondence at the loci level. In terms of unique loci, 5,233 genes annotated by Ensembl (including 3,950 lncRNAs and 828 PCGs) had no matching gene model in Refseq, while 5,002 genes were specific to Refseq (including 3,024 lncRNAs and 1,363 PCGs) (Figure 7B). Note that, 148 miRNAs were annotated exclusively by Ensembl, whereas 260 tRNAs were found only in Refseq and were not previously represented in the Ensembl dataset. Gene fusion events were also detected during the merging process. A total of 460 fusion cases were identified, of which more than 95% involved the combination of exactly two genes. For example, 67 Ensembl genes were found to encompass multiple Refseq genes (n=136), and 325 Refseq genes incorporated multiple Ensembl gene models (n=704). For example, the TTN gene (LOC107316616), known to be one of the longest (Bang et al. 2001), was identified as a unique locus in Refseq but was divided into 14 loci in Ensembl (Sup. Figure 1). This gene was merged with the enhanced annotation and seems to correspond better to its well-known ortholog in the human genome. For genes with unambiguous one-to-one matches, more than 99.5% concordance in gene biotype was observed. For example, only 14 genes annotated as PCGs in Refseq were classified as lncRNAs in Ensembl, and similar biotype consistency was noted among merged gene models. At the transcript level, biotype consistency was fully maintained, as all original transcripts were retained in the merged annotation, or their source identifiers were preserved when exon chains were identical.

### Transfer of the Enhanced Annotation to the *Cjap1.hap1* Assembly

The enhanced annotation was transferred to the new *Coturnix japonica* genome assembly (Cjap1.hap1) using the LiftOff pipeline. A total of 25,038 genes and 84,405 transcripts were successfully mapped, corresponding to 96% and 97% of the gene and transcript models from the merged annotation, respectively. Among the transferred gene models, 16,122 PCGs and 7,822 lncRNAs were identified (Figure 7A).

The quality of the lift-over was evaluated based on gene model coverage and sequence identity (Figure 7D). Overall, 95% of lifted genes displayed a coverage ≥90% and a sequence identity ≥80%. As expected, based on their evolutionary constraints, PCGs exhibited higher transfer fidelity, with 95% of models showing a coverage ≥96%. In contrast, 90% of lncRNA models achieved a minimum of 90% coverage. Similar patterns were observed for sequence identity, where PCGs consistently exhibited higher conservation than lncRNAs.

To characterize the 1,064 gene models that were not successfully lifted—comprising 639 PCGs and 349 lncRNAs—a nucleotide sequence similarity search was performed using BLASTn against the new *Cjap1.hap1* genome assembly. For benchmarking, genes that had been successfully lifted were also included as a control. Among the lifted genes, 98.7% (24,723/25,037) produced significant BLASTn hits, confirming their mappability. In contrast, only 48.8% (519/1,064) of the previously unmapped gene models returned significant hits, underscoring the challenges associated with lifting these sequences to the new assembly (Figure 7C). Furthermore, the 519 unmapped genes with BLASTn matches exhibited significantly higher E-values (log10 transformation; -194 vs. -126; t-test; pval<10^-16^) compared to those from the previously mapped set, indicating weaker sequence similarity and suggesting possible assembly-related or annotation-specific discrepancies. Conversely, for the 314 gene models that had been lifted by LiftOff but did not produce any BLASTn hit, a significantly lower coverage (0.98 vs. 0.50; t-test; pval<10^-16^) was observed compared to the rest of the lifted genes, indicating a partial or fragmented mapping that may have affected their alignment detectability.

### Expression quantification using the transfer-enhanced annotation

Gene expression quantification was carried out using the enriched annotation lifted onto the Cjap1.hap1 assembly, across seven samples corresponding to seven distinct adult tissues. As expected, a higher number of PCGs were detected as expressed (defined as ≥0.1 TPM) relative to lncRNAs, with median values of 13,474 and 1,597 genes per tissue, respectively—corresponding to 84% and 20% of the total annotated gene sets. When expressed, PCGs exhibited significantly higher expression levels than lncRNAs, with a mean of median expression by tissue of 11.02 TPM versus 0.41 TPM (Figure 8A).

**Figure 8.**
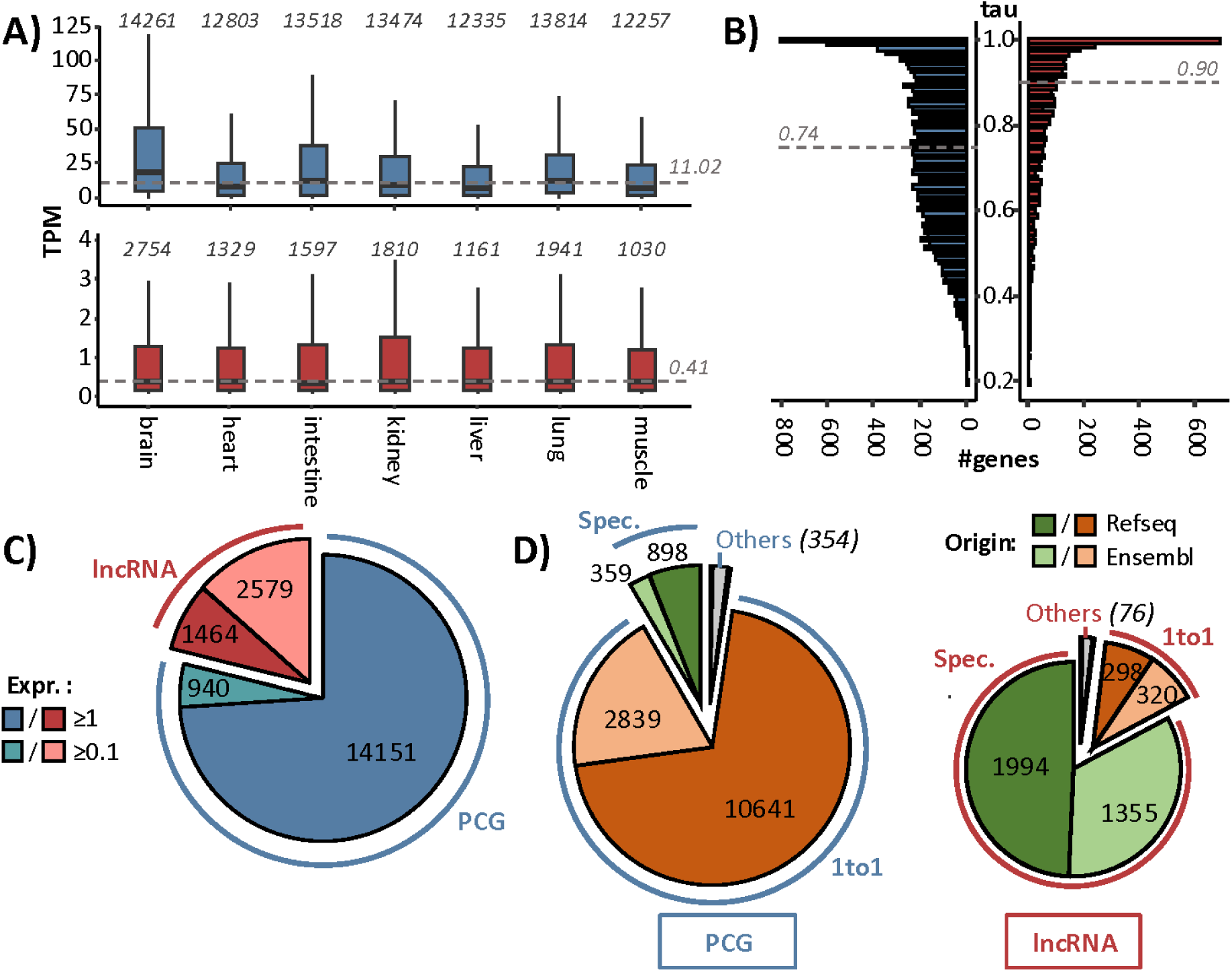
Gene expression profiles from the enhanced annotation on the *Cjap1.hap1* assembly. **A,** Expression distribution (TPM) of protein-coding genes (blue) and long non-coding RNAs (lncRNAs, red) expressed (≥0.1 TPM) across seven tissues. Numbers above each boxplot indicate the number of expressed genes per tissue. The grey line denotes the median expression level across all tissues. **B,** Tissue specificity, measured using the τ (tau) metric, for each gene biotype. The grey line represents the median τ value. **C,** Number of genes expressed in at least one of the seven tissues, grouped by biotype and expression threshold. **D,** Number of expressed genes stratified by annotation origin.

Expression specificity, measured using the tissue-specificity index (τ), revealed that lncRNAs tend to exhibit more tissue-restricted expression patterns. The median τ value reached 0.90 for lncRNAs, in contrast to 0.74 for PCGs, confirming the broader expression of PCGs across tissues and the more specific expression of lncRNAs (Figure 8B). Finally, when considering expression across all the seven tissues, 15,091 PCGs (94%) and 4,043 lncRNAs (52%) were found to be expressed in at least one tissue (Figure 8C).

The origin of the expressed genes was also examined. Among genes shared between the Ensembl and Refseq annotations, 14,247 out of 15,393 (93%) were expressed, suggesting strong support for these consensus annotations (Figure 8D). In contrast, 2,910 out of 5,002 genes unique to Refseq (58%), and 1,844 out of 5,233 Ensembl-specific genes (35%) were detected as expressed, reflecting both the complementary nature and potential variability in annotation source specificity.

## Discussion

Here we present two new assemblies of the quail genome, with a better contiguity than the current version (Contig N50 Cjap1.hap1 = 6.8 Mb) and a good completeness (Cjap1.hap1 BUSCO score - aves_odb10 lineage = 97.3). While the BUSCO scores between both assemblies are quite similar (96.5 for CoJa2.1), the Contig N50 is 8.6 times higher in the new assembly. The length of the obtained assembly (1.19) is closer than CoJa2.1 (0.93 Mb) to the expected genome size for this species (1.29 to 1.41 Mb, (Gregory 2025)). Notably, the assembly extends previously truncated chromosome ends, with 31 telomeres identified, while no telomere was detected in CoJa2.1.

The high degree of collinearity observed for both macrochromosomes and the Z chromosome demonstrates the robustness of the updated assembly at the chromosome scale. Notably, the anchoring of approximately 75% of previously unlocalized contigs to their corresponding chromosomes, especially the complete placement of contigs from eight chromosomes, marks a significant advance in genome completeness. This improvement is exemplified by chromosome 1, which not only increases in size by 2.3% but also integrates both previously unlocalized and unplaced contigs, reflecting enhanced resolution. Nevertheless, the cumulative contribution of these newly placed sequences might be smaller than their earlier unplaced total, indicating that previous contigs likely contained redundant or repetitive sequences, and that part of the improvement also reflects a consolidation rather than simply an increase in sequence content. Furthermore, alignment with the chicken genome enabled the provisional identification of additional microchromosomes, suggesting that the Cjap1.hap1 assembly captures genomic features that were previously unresolved. While these observations provide compelling candidates for unresolved microchromosomes, their assignment remains provisional. Further manual curation and complementary data will be necessary to fully characterize and annotate the full repertoire of microchromosomes in Cjap1.hap1. As already highlighted (Hillier et al. 2004; Morris et al. 2020), the quail and chicken genomes, which diverged around 35–40 million years ago, retain the conserved macro- and microchromosome organization that constitutes a signature of avian genomes. Although we had sequenced a male, the results obtained did not allow us to assemble the W chromosome.

Because the quality of a genome assembly also depends on the assembler, the same long read data sets has been used to compare assemblers in (Birbes et al. 2025). Hifiasm did not perform best on metrics such as NG50 and LG50. ONT reads assembled with NextDeNovo produced an assembly with a 13.00 Mb NG50 and a 21 contigs LG50 compared to 4.05 Mb and 84 contigs for the same metrics with hifiasm using only HiFi reads. These metrics have been slightly improved by adding the ONT long reads to the HiFi reads while running hifiasm for Cjap1.hap1.1 but are only half as good as ONT only. The possible cause of this are AG rich genomic regions which are poorly or not represented in HiFi reads as mentioned in (Rabanal et al. 2022). Because no HiFi reads are produced for these regions, they are gapped in the assembly.

With the assembly quality assessed, the subsequent step consisted in annotating the genome to enable functional analyses.

Rather than generating annotations entirely de novo, we adopted a strategy of leveraging high-quality existing resources. Both Refseq and Ensembl annotations integrate evidence from ab initio predictions and RNA-seq data, and are curated through rigorous pipelines (Goldfarb et al. 2025). Considering current resource limitations—including lack of deep transcriptomic coverage across tissues in quail—this approach was deemed both efficient and robust. While long-read RNA-seq or capture-based methods would further improve annotation quality, especially to identify novel lncRNAs or to improve isoform discovery for PCGs, such efforts are resource-intensive due to their requirement for extensive tissue sampling and computational processing (Pardo-Palacios et al. 2024; Carbonell-Sala et al. 2024). For annotation transfer, we employed LiftOff (Shumate and Salzberg 2021), a gene-aware liftover tool that maximizes the preservation of exon-intron structure and transcript models, in contrast to coordinate-based tools such as UCSC LiftOver that rely solely on chain file alignment (Navarro Gonzalez et al. 2021; Genovese et al. 2024). Although we observed high concordance between LiftOff and BLASTN-based projection, LiftOff showed better agreement at the transcript level, justifying its use as the primary tool. To preserve isoform granularity, transcript definitions were retained at the exon level in the final merged annotation. Although intron-chain–based simplification could have reduced annotation complexity, such reduction (∼8% fewer transcripts) risked losing biologically relevant isoforms with alternative UTRs.

While the enhanced annotation provides a more comprehensive and inclusive representation of the *Coturnix japonica* genome, a limited number of ambiguous cases remain, particularly regarding gene boundary definitions and biotype classification. These rare discrepancies underscore the value of integrating multiple annotation sources, as already emphasized in different comparative annotation works (Degalez et al. 2024; Kaur et al. 2024). By preserving all transcript-level information during the integration process, most of these conflicts were mitigated. Indeed, even in instances where gene coordinates were difficult to reconcile, the associated transcripts were retained unaltered, maintaining their original structure. This transcript-centric strategy aligns with the growing emphasis on isoform resolution afforded by long-read sequencing technologies, which enable accurate transcript-level reconstruction.

Biotypes specific to a single source, such as tRNAs from Refseq or miRNAs from Ensembl, were retained despite their underrepresentation in standard RNA-seq datasets focusing mainly on long transcripts. Nevertheless, inclusion of these biotypes enhances the utility of the annotation for downstream analyses, including studies on small RNA pathways and comparative genomics.

Our expression analysis further supports the biological relevance of the enhanced annotation. Notably, 94% of PCGs and 52% of lncRNAs were expressed in at least one of the seven tissues sampled. Consistent with prior studies (Derrien et al. 2012; Guan et al. 2025; Degalez et al. 2024), PCGs were broadly expressed across tissues and at higher levels than lncRNAs, which were more tissue-specific and typically lower in abundance. Despite the limited tissue set, these results underscore the value of incorporating lncRNAs and other non-coding biotypes in genome annotation. Future studies expanding tissue coverage and employing long-read transcriptomics will help refine tissue-specific expression estimates and uncover additional functional lncRNAs. Importantly, while most expressed PCGs were shared between Refseq and Ensembl annotations, the majority of expressed lncRNAs were specific to a single source. This observation reflects both the complementary nature of annotation pipelines and the variability inherent in lncRNA prediction—highlighting the need for continued refinement of non-coding gene annotation across species.

The newly assembled and annotated quail genome offers a high-quality resource that will facilitate future research in avian biology and agriculture.

## Acknowledgements

We are grateful to the entire staff of the INRAE experimental unit PEAT, Nouzilly, https://doi.org/10.15454/1.5572326250887292E12) for their excellent animal care. This work was performed in collaboration with the GeT core facility, Toulouse, France (GeT, https://doi.org/10.15454/1.5572370921303193E12).

## Data, scripts, code, and supplementary information availability

Data are available online with the accession numbers ERR11528759 and ERR11528760 for HiFi, ERR11591487 and ERR11591488 for ONT and ERR13583727 and ERR13583728 for Hi-C (ENA database, https://www.ebi.ac.uk/ena/browser).

The genome assembly is available at European Nucleotide Archive (https://www.ebi.ac.uk/ena/) as two haplotypes: accession numbers GCA_978018805.1 and GCA_977970185.1).

Scripts and code are available online: https://zenodo.org/records/17122626 (doi:10.5281/zenodo.17122625).

## Supplementary Figures

**Sup. Figure 1.**
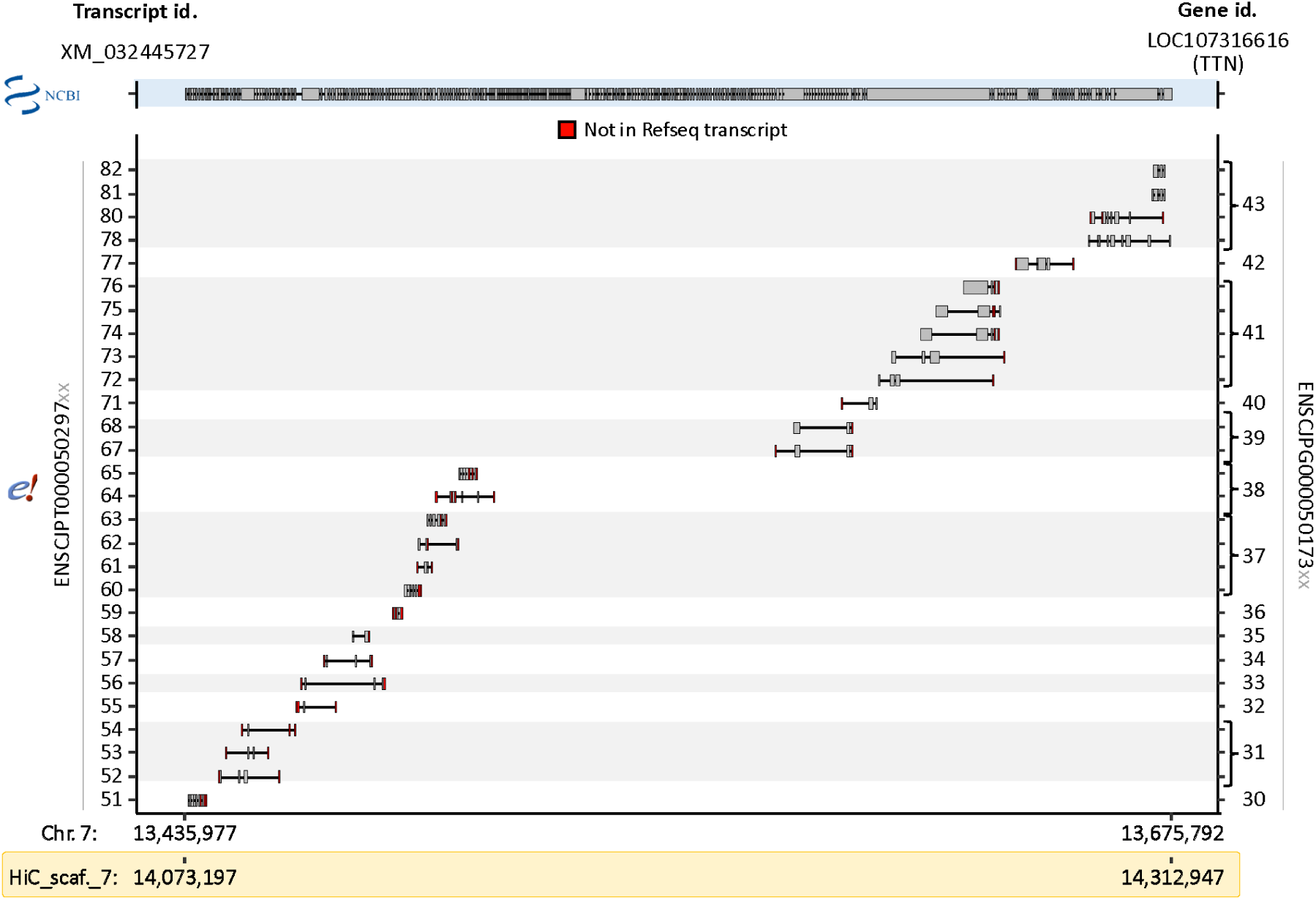
Comparison of the TTN gene identified in Refseq and the associated truncated models in Ensembl.

## References

Alonge, Michael, Ludivine Lebeigle, Melanie Kirsche, et al. 2022. « Automated assembly scaffolding using RagTag elevates a new tomato system for high-throughput genome editing ». Genome Biology 23 (1): 258. 10.1186/s13059-022-02823-7.

Balthazart, J., M. Baillien, T. D. Charlier, C. A. Cornil, et G. F. Ball. 2003. « The neuroendocrinology of reproductive behavior in Japanese quail ». *Domestic Animal Endocrinology*, Reproductive Physiology: Proceedings of a workshop held at the Institute for Animal Science Mariensee, Germany, vol. 25 (1): 69-82. 10.1016/S0739-7240(03)00046-8.

Bang, Marie-Louise, Thomas Centner, Friderike Fornoff, et al. 2001. « The Complete Gene Sequence of Titin, Expression of an Unusual ≈700-kDa Titin Isoform, and Its Interaction With Obscurin Identify a Novel Z-Line to I-Band Linking System ». Circulation Research 89 (11): 1065-72. 10.1161/hh2301.100981.

Barzilai-Tutsch, Hila, Valerie Morin, Gauthier Toulouse, et al. 2022. « Transgenic quails reveal dynamic TCF/β-catenin signaling during avian embryonic development ». eLife 11 (juillet): e72098. 10.7554/eLife.72098.

Birbes, Clément, Andreea Dréau, Denis Milan, et al. 2025. « Benchmarking Long-Read Genome Assemblers for Three Sequencing Protocols and Three Agricultural Species ». Prépublication, bioRxiv, avril 22. 10.1101/2025.02.14.638238.

Carbonell-Sala, Sílvia, Tamara Perteghella, Julien Lagarde, et al. 2024. « CapTrap-Seq: A Platform-Agnostic and Quantitative Approach for High-Fidelity Full-Length RNA Sequencing ». Nature Communications 15 (1): 5278. 10.1038/s41467-024-49523-3.

Challis, Richard, Edward Richards, Jeena Rajan, Guy Cochrane, et Mark Blaxter. 2020. « BlobToolKit – Interactive Quality Assessment of Genome Assemblies ». G3 Genes|Genomes|Genetics 10 (4): 1361-74. 10.1534/g3.119.400908.

Chazara, Olympe, Francis Minvielle, Denis Roux, et al. 2010. « Evidence for Introgressive Hybridization of Wild Common Quail (Coturnix Coturnix) by Domesticated Japanese Quail (Coturnix Japonica) in France ». Conservation Genetics 11 (3): 1051-62. 10.1007/s10592-009-9951-8.

Cheng, Haoyu, Gregory T. Concepcion, Xiaowen Feng, Haowen Zhang, et Heng Li. 2021. « Haplotype-Resolved de Novo Assembly Using Phased Assembly Graphs with Hifiasm ». Nature Methods 18 (2): 170-75. 10.1038/s41592-020-01056-5.

Dainat, Jacques, Robrecht Cannoodt, André Soares, et al. 2025. NBISweden/AGAT: AGAT v1.5.1. Zenodo, released juillet 22. 10.5281/zenodo.16317950.

Degalez, Fabien, Mathieu Charles, Sylvain Foissac, et al. 2024. « Enriched Atlas of lncRNA and Protein-Coding Genes for the GRCg7b Chicken Assembly and Its Functional Annotation across 47 Tissues ». Scientific Reports 14 (1): 6588. 10.1038/s41598-024-56705-y.

Derrien, Thomas, Rory Johnson, Giovanni Bussotti, et al. 2012. « The GENCODE v7 Catalog of Human Long Noncoding RNAs: Analysis of Their Gene Structure, Evolution, and Expression ». Genome Research 22 (9): 1775-89. 10.1101/gr.132159.111.

Du, Kang, Matthias Stöck, Susanne Kneitz, et al. 2020. « The Sterlet Sturgeon Genome Sequence and the Mechanisms of Segmental Rediploidization ». Nature Ecology & Evolution 4 (6): 841-52. 10.1038/s41559-020-1166-x.

Dudchenko, Olga, Sanjit S. Batra, Arina D. Omer, et al. 2017. « De novo assembly of the Aedes aegypti genome using Hi-C yields chromosome-length scaffolds ». Science 356 (6333): 92-95. 10.1126/science.aal3327.

Durand, Neva C., James T. Robinson, Muhammad S. Shamim, et al. 2016. « Juicebox Provides a Visualization System for Hi-C Contact Maps with Unlimited Zoom ». Cell Systems 3 (1): 99-101. 10.1016/j.cels.2015.07.012.

Durand, Neva C., Muhammad S. Shamim, Ido Machol, et al. 2016. « Juicer Provides a One-Click System for Analyzing Loop-Resolution Hi-C Experiments ». Cell Systems 3 (1): 95-98. 10.1016/j.cels.2016.07.002.

Foissac, Sylvain, Sarah Djebali, Kylie Munyard, et al. 2019. « Multi-species annotation of transcriptome and chromatin structure in domesticated animals ». BMC Biology 17 (décembre): 108. 10.1186/s12915-019-0726-5.

Gel, Bernat, et Eduard Serra. 2017. « karyoploteR: an R/Bioconductor package to plot customizable genomes displaying arbitrary data ». Bioinformatics 33 (19): 3088-90. 10.1093/bioinformatics/btx346.

Genovese, Giulio, Nicole B Rockweiler, Bryan R Gorman, et al. 2024. « BCFtools/liftover: an accurate and comprehensive tool to convert genetic variants across genome assemblies ». Bioinformatics 40 (2): btae038. 10.1093/bioinformatics/btae038.

Goldfarb, Tamara, Vamsi K Kodali, Shashikant Pujar, et al. 2025. « NCBI RefSeq: reference sequence standards through 25 years of curation and annotation ». Nucleic Acids Research 53 (D1): D243-57. 10.1093/nar/gkae1038.

Gregory, T.R. 2025. « Animal Genome Size Database. » http://www.genomesize.com.

Guan, Dailu, Zhonghao Bai, Xiaoning Zhu, et al. 2025. « Genetic Regulation of Gene Expression across Multiple Tissues in Chickens ». Nature Genetics 57 (5): 1298-308. 10.1038/s41588-025-02155-9.

Hillier, LaDeana W., Webb Miller, Ewan Birney, et al. 2004. « Sequence and Comparative Analysis of the Chicken Genome Provide Unique Perspectives on Vertebrate Evolution ». Nature 432 (7018): 695-716. 10.1038/nature03154.

Huss, David, Bertrand Benazeraf, Allison Wallingford, et al. 2015. « A Transgenic Quail Model That Enables Dynamic Imaging of Amniote Embryogenesis ». *Development (Cambridge*, England) 142 (16): 2850-59. 10.1242/dev.121392.

Kaur, Gazaldeep, Tamara Perteghella, Sílvia Carbonell-Sala, et al. 2024. « GENCODE: Massively Expanding the lncRNA Catalog through Capture Long-Read RNA Sequencing ». Prépublication, bioRxiv, octobre 31. 10.1101/2024.10.29.620654.

Kayang, Boniface B., Valérie Fillon, Miho Inoue-Murayama, et al. 2006. « Integrated maps in quail (Coturnix japonica) confirm the high degree of synteny conservation with chicken (Gallus gallus) despite 35 million years of divergence ». BMC Genomics 7 (1): 101. 10.1186/1471-2164-7-101.

Le Douarin, Nicole. 1973. « A biological cell labeling technique and its use in experimental embryology ». Developmental Biology 30 (1): 217-22. 10.1016/0012-1606(73)90061-4.

Le Douarin, Nicole. 2005. « The Nogent Institute - 50 Years of Embryology ». The International Journal of Developmental Biology 49 (2-3): 85-103. 10.1387/ijdb.041952nl.

Leroux, Sophie, David Gourichon, Christine Leterrier, et al. 2017. « Embryonic Environment and Transgenerational Effects in Quail ». Genetics, Selection, Evolution: GSE 49 (1): 14. 10.1186/s12711-017-0292-7.

Li, Heng. 2018. « Minimap2: pairwise alignment for nucleotide sequences ». Bioinformatics 34 (18): 3094-100. 10.1093/bioinformatics/bty191.

Lin, C.-Y., C.-H. Ho, Y.-H. Hsieh, et T. Kikuchi. 2002. « Adeno-Associated Virus-Mediated Transfer of Human Acid Maltase Gene Results in a Transient Reduction of Glycogen Accumulation in Muscle of Japanese Quail with Acid Maltase Deficiency ». Gene Therapy 9 (9): 554-63. 10.1038/sj.gt.3301672.

Lukanov, H. 2019. « Domestic quail (Coturnix japonica domestica), is there such farm animal? » World’s Poultry Science Journal 75 (4): 547-58. 10.1017/S0043933919000631.

Manni, Mosè, Matthew R Berkeley, Mathieu Seppey, Felipe A Simão, et Evgeny M Zdobnov. 2021. « BUSCO Update: Novel and Streamlined Workflows along with Broader and Deeper Phylogenetic Coverage for Scoring of Eukaryotic, Prokaryotic, and Viral Genomes ». Molecular Biology and Evolution 38 (10): 4647-54. 10.1093/molbev/msab199.

Mignon-Grasteau, S, O Roussot, C Delaby, et al. 2003. « Factorial correspondence analysis of fear-related behaviour traits in Japanese quail ». Behavioural Processes 61 (1): 69-75. 10.1016/S0376-6357(02)00162-6.

Mills, A. D., et J. M. Faure. 1991. « Divergent Selection for Duration of Tonic Immobility and Social Reinstatement Behavior in Japanese Quail (Coturnix Coturnix Japonica) Chicks ». Journal of Comparative Psychology (Washington, D.C.: 1983) 105 (1): 25-38. 10.1037/0735-7036.105.1.25.

Morris, Katrina M., Matthew M. Hindle, Simon Boitard, et al. 2020. « The quail genome: insights into social behaviour, seasonal biology and infectious disease response ». BMC Biology 18 (février): 14. 10.1186/s12915-020-0743-4.

Navarro Gonzalez, Jairo, Ann S Zweig, Matthew L Speir, et al. 2021. « The UCSC Genome Browser database: 2021 update ». Nucleic Acids Research 49 (D1): D1046-57. 10.1093/nar/gkaa1070.

Pardo-Palacios, Francisco J., Dingjie Wang, Fairlie Reese, et al. 2024. « Systematic Assessment of Long-Read RNA-Seq Methods for Transcript Identification and Quantification ». Nature Methods 21 (7): 1349-63. 10.1038/s41592-024-02298-3.

Patel, Harshil, Phil Ewels, Jonathan Manning, et al. 2024. *nf-core/rnaseq: nf-core/rnaseq v3.16.0 - Fire Ferret*. Zenodo, released octobre 2. 10.5281/zenodo.13882081.

Peona, Valentina, Matthias H. Weissensteiner, et Alexander Suh. 2018. « How Complete Are “Complete” Genome Assemblies?-An Avian Perspective ». Molecular Ecology Resources 18 (6): 1188-95. 10.1111/1755-0998.12933.

Rabanal, Fernando A, Maike Gräff, Christa Lanz, et al. 2022. « Pushing the limits of HiFi assemblies reveals centromere diversity between two Arabidopsis thaliana genomes ». Nucleic Acids Research 50 (21): 12309-27. 10.1093/nar/gkac1115.

Roussot, Odile, Katia Feve, Florence Plisson-Petit, et al. 2003. « AFLP Linkage Map of the Japanese Quail Coturnix Japonica ». Genetics, Selection, Evolution: GSE 35 (5): 559-72. 10.1186/1297-9686-35-6-559.

Sáez, David, Fernando Spina, Antoni Margalida, Lorenzo Serra, Stefano Volponi, et Jesús Nadal. 2023. « Reconstructing Migratory Network Nodes to Improve Environmental Management and Conservation Decisions: A Case Study of the Common Quail Coturnix Coturnix as a Biosensor ». The Science of the Total Environment 893 (octobre): 164913. 10.1016/j.scitotenv.2023.164913.

Sanchez-Donoso, Ines, Pablo Antonio Morales-Rodriguez, Manel Puigcerver, José Ramón Caballero de la Calle, Carleso Vilà, et José Domingo Rodríguez-Teijeiro. 2016. « Postcopulatory Sexual Selection Favors Fertilization Success of Restocking Hybrid Quails over Native Common Quails (Coturnix Coturnix) ». Journal of Ornithology 157 (1): 33-42. 10.1007/s10336-015-1242-1.

Serralbo, Olivier, David Salgado, Nadège Véron, et al. 2020. « Transgenesis and web resources in quail ». eLife 9: e56312. 10.7554/eLife.56312.

Shumate, Alaina, et Steven L Salzberg. 2021. « Liftoff: accurate mapping of gene annotations ». Bioinformatics 37 (12): 1639-43. 10.1093/bioinformatics/btaa1016.

Smith, Jacqueline, James M. Alfieri, Nick Anthony, et al. 2023. « Fourth Report on Chicken Genes and Chromosomes 2022 ». Cytogenetic and Genome Research 162 (8-9): 405-528. 10.1159/000529376.

Talenti, Andrea, et James Prendergast. 2021. « nf-LO: A Scalable, Containerized Workflow for Genome-to-Genome Lift Over ». Genome Biology and Evolution 13 (9): evab183. 10.1093/gbe/evab183.

Terzian, Paul, Céline Vandecasteele, Joanna Lledo, et al. 2025. « Pig and Quail CpG Methylation Datasets from Short and Long Read Sequencing Technologies ». Scientific Data 12 (1): 556. 10.1038/s41597-025-04769-4.

Uliano-Silva, Marcela, João Gabriel R. N. Ferreira, Ksenia Krasheninnikova, et al. 2023. « MitoHiFi: a python pipeline for mitochondrial genome assembly from PacBio high fidelity reads ». BMC Bioinformatics 24 (1): 288. 10.1186/s12859-023-05385-y.

Vitorino Carvalho, Anaïs, Christelle Hennequet-Antier, Romuald Rouger, et al. 2023. « Thermal Conditioning of Quail Embryos Has Transgenerational and Reversible Long-Term Effects ». Journal of Animal Science and Biotechnology 14 (1): 124. 10.1186/s40104-023-00924-2.

Yanai, Itai, Hila Benjamin, Michael Shmoish, et al. 2005. « Genome-wide midrange transcription profiles reveal expression level relationships in human tissue specification ». Bioinformatics 21 (5): 650-59. 10.1093/bioinformatics/bti042.

